# miRNAtools: advanced training using the miRNA web of knowledge

**DOI:** 10.1101/170126

**Authors:** Francisco J. Enguita

## Abstract

miRNAs are small non-coding RNAs, that act as negative regulators of the genomic output. Their intrinsic importance within cell biology and human disease is well known. Their mechanism of action based on the base pairing binding to their cognate targets, have helped the development of many computer applications for the prediction of miRNA target recognition. More recently, many other computer applications have appeared with the objective of studying miRNA roles in many contexts, trying to dissect and predict their functions in a specific biological process. Learning about miRNA function needs a practical training in the use of specific computer and web-based applications that are complementary to the wet-lab studies. In the last seven years we have been involved in the organization of advanced training courses about the *in silico* functional analysis of miRNAs and non-coding RNAs, for students ranging from the postgraduate to the post-doctoral level. In order to guide the learning process about miRNAs, we have created miRNAtools (http://mirnatools.eu), a web repository of miRNA tools and tutorials. This page is a compilation of tools to analyze miRNAs and their regulatory action that intends to collect and organize the information that is dispersed in the web. The miRNAtools webpage is completed by a collection of tutorials that can be used by students and tutors engaged in advanced training courses. The tutorials follow the rationale of the analysis of the function of selected miRNAs, starting from their nomenclature and genomic localization and finishing by assessing their involvement in specific cellular functions.

**DESCRIPTION OF THE AUTHOR:** Francisco J. Enguita is assistant professor at the Faculty of Medicine, University of Lisbon and senior investigator at the Instituto de Medicina Molecular within the same University.

**KEY POINTS:**

1. - We have designed a web-page, mIRNAtools3 (http://mirnatools.eu) specifically devoted for advanced teaching purposes within the miRNA field.
2. - The webpage constitutes also a repository of web-tools for the analysis of miRNA function in several contexts.
3. - The webpage contains tutorials that cover many aspects related with miRNAs including nomenclature, target prediction and validation, and functional analysis.

## 1.- Introduction

Micro-RNAs (miRNAs) are small non-coding RNAs (ncRNAs), firstly described in the early 90’s by Victor Ambros and Gary Ruvkun research groups [1, 2]. These tiny ncRNAs associated with specific proteins act as negative post-transcriptional regulators by binding to the 3’-UTR of mRNA transcripts. The miRNA binding to the 3’-UTR of a mRNAs is based on their complementarity and ensured by the Watson-Crick base pairing rules. In animals the miRNA is only partially complementary to its mRNA target, typically involving the nucleotides 2 to 9 of its 5’ end (seed sequence). However in plants, the complementarity of the miRNA and its target is typically higher than 90%. These different pairing rules in animals and plants are translated in different immediate regulatory effects. Whereas in animals the partial complementarity produces a translational repression as a main regulatory effect [3], the almost complete complementarity between miRNAs and their targets observed in plants ensures a degradation of the target mRNA as a primary consequence [4, 5].

MiRNAs are global regulators of cellular pathways, constituting a layer of regulation which is superimposed to the classical transcriptional regulators. The small size of miRNAs is responsible for their regulatory promiscuity: a single miRNA can regulate hundreds of different miRNA transcripts and at the same time, a given mRNA transcript can be targeted by several miRNAs. However, the regulatory effect exerted by a miRNA is only possible if the miRNA and its cognate target are present simultaneously in the same biological environment. This is the subjacent reason below the variety of regulatory effects of the same miRNA when we consider a different cellular context.

Since the discovery of miRNAs many bioinformatics tools have been developed with the objective of study their physiological roles [6–8]. Among the bioinformatics applications related to miRNAs, target prediction tools are the most important and the basis for the majority of the existing tools for miRNA functional analysis [9–11]. Target prediction algorithms have evolved along time from the classical ones that only took into account the base complementarity roles between miRNAs and their targets [3, 6], to the modern ones based on deep learning and neural network algorithms [12, 13]. Other computer applications take advantage of the target prediction algorithms to infer physiological roles for miRNAs [14–16]. The evolution of scientific protocols has also dramatically changed the needs for computer applications related to miRNAs. For instance, some of the original applications designed for the prediction of miRNA genes across genomes have becalmed obsolete with the introduction of the next-generation sequencing techniques [17, 18]. In fact, the web landscape is densely populated with web tools designed for miRNA analysis, that in many occasions are not so easy to find and to use.

## 2.- The need for advanced teaching resources for miRNA functional analysis

During the last six years we have been involved in advanced teaching initiatives for students in biomedical sciences ranging from the post-graduate to the post-doctoral level within the field of miRNAs and other non-coding RNAs (ncRNAs). We have organized lecture and practical courses and workshops in our Institution and abroad to introduce the field of miRNAs in a practical manner using bioinformatics tools. From the very beginning it was evident for us that the students were facing the problem of the existence of myriad of bioinformatics tools dispersed in the web that sometimes were not easy to use and interpret. Moreover, the fast evolution of laboratory analytical techniques and computer software made some of these tools to become obsolete very quickly. Some excellent compilations of miRNA web-tools such as Tools4miRs [19], miRandb [20] and miRtoolsGallery (http://mirtoolsgallery.org/miRToolsGallery) have been designed with the main objective of systematize, classify and organize all the available web-tools within the field of miRNAs. However, there was a clear need for specialized teaching material devoted to the study of miRNAs and other ncRNAs that could be employed for self-teaching and also as conducting script for advanced courses. For all the exposed reasons, we created miRNAtools (http://mirnatools.eu), a repository of web tools devoted to the study of miRNAs which is complemented by specialized tutorials for assisted or self-learning. The main difference between miRNAtools3 and all other existing repositories of web-tools is its specific design for advanced teaching.

## 3.- Structure of miRNAtools3 webpage

MiRNAtools webpage (http://mirnatools.eu) has evolved from the preliminary beta version launched in 2010 to the new miRNAtools3 which has been made available on the web in May 2017. MiRNAtools3 is built in HTML 5.0, and complemented with CSS and Javascript aids that allow a rapid and easy navigation between sections. The overall flexible web design ensures a rapid update of sections and the inclusion of new topics, tutorials and applications without changing the general outlook.

MiRNAtools3 webpage contains two dependence levels (Figure 1). The first one is currently structured in six main sections: databases, targets, NGS tools, *In silico* tools, Plant miRNAs and Tutorials. The “Databases” section contains two sub-sections compiling “General purpose databases”, compiling links to miRNA-related databases for general use, and “Specialized databases” that contain databases designed for studying specific sub-fields of the miRNA world such as circulating miRNAs, evolution and function of miRNAs and disease-related databases.

**Figure 1:**
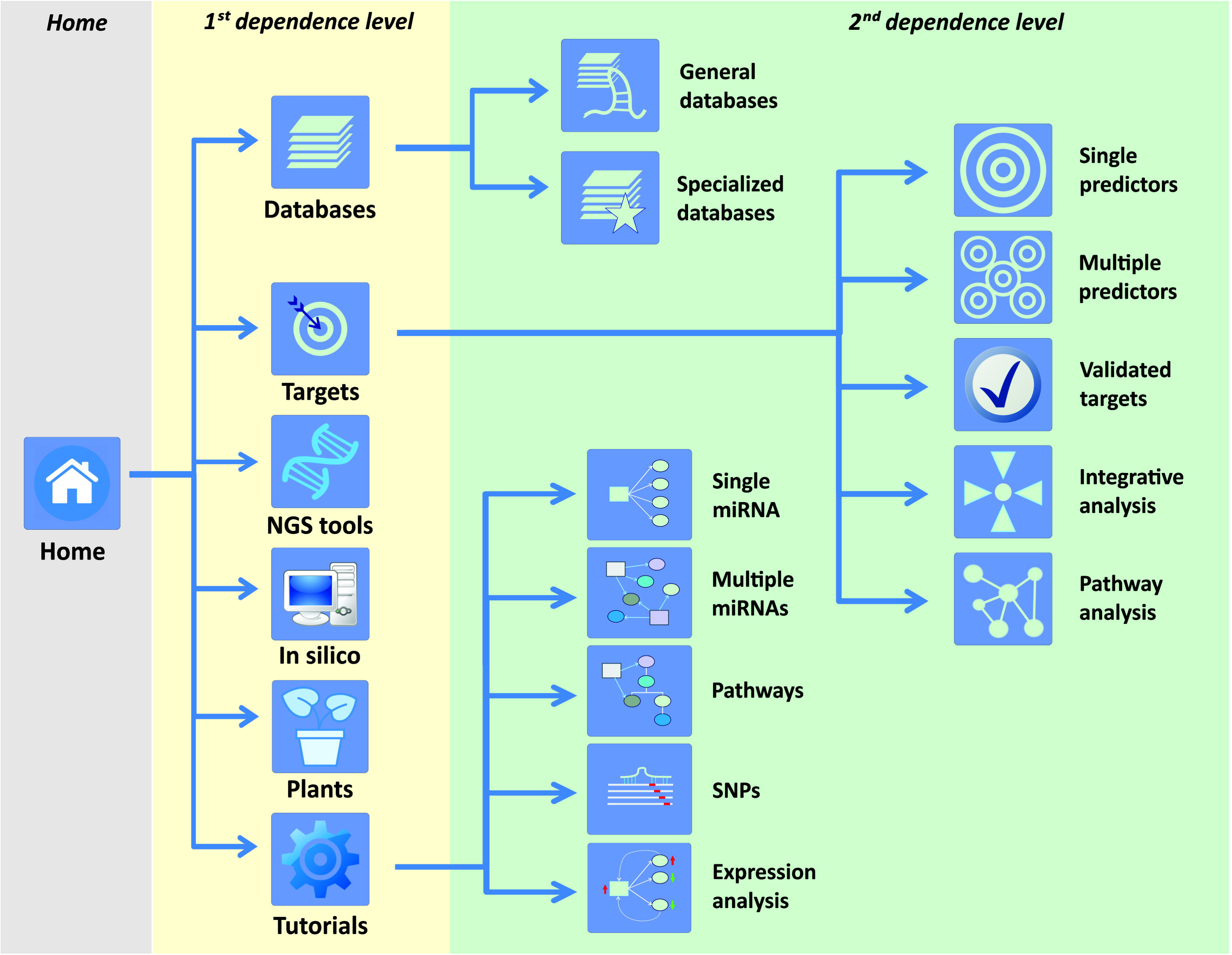
navigation structure of miRNAtools3 webpage showing the two main dependence levels. Navigation across sections and topics can be achieved by direct pointing over the respective icons or by using the navigation menu integrated in each sub-page.

The “Targets” first dependence level is the most complex, and includes five sub-sections: single predictors, multiple predictors, validated targets, integrative analysis and pathway analysis. The “Single predictors” sub-section is devoted to compile web-applications for miRNA target prediction. Every link to an external web-page is accompanied by a personal assessment of the advantages and disadvantages of each algorithm and the corresponding link to a Pubmed reference. The “Multiple predictors” second dependence level includes web-based applications able to interrogate multiple target predictors at the same time. These applications are especially powerful since they are able to generate a stronger and more reliable target prediction than the individual algorithms considered alone. The “Validated targets” sub-section includes web-tools and databases for the retrieval of validated miRNA targets. The last two-subsections within the “Targets” first dependence level are devoted to the functional analysis of miRNA action. The “Integrative analysis” contains web-tools designed for the integration of transcriptomic data and miRNA expression, and the “Pathway analysis” sub-section is composed by bionformatics tools that infer the action of one or several miRNAs over cell metabolic pathways. Every item within sections and sub-sections contains a personal overview and description and the reference to its publication. The first dependence level of miRNAtools3 is completed by three additional independent sections covering tools related with miRNA target analysis in plants, *in silico* tools for the prediction of novel miRNA genes in new genomes, tools for next-generation sequencing analysis of miRNA expression, and tutorials.

## 4.- Tutorials within miRNAtools3

The tutorial section is designed for self-teaching, but also to serve as a main guiding script for specialized courses or workshops. This section is divided into proposed scenarios that introduce a biological example involving a particular miRNA or groups of miRNAs and guide the user through different web applications to obtain relevant information about the biological problem. All the scenarios are explained by using screenshots of the selected web-tools in order to guide the student about the inputs and outputs of every application. The specific applications and topics covered in each tutorial are depicted on Figure 2.

**Figure 2:**
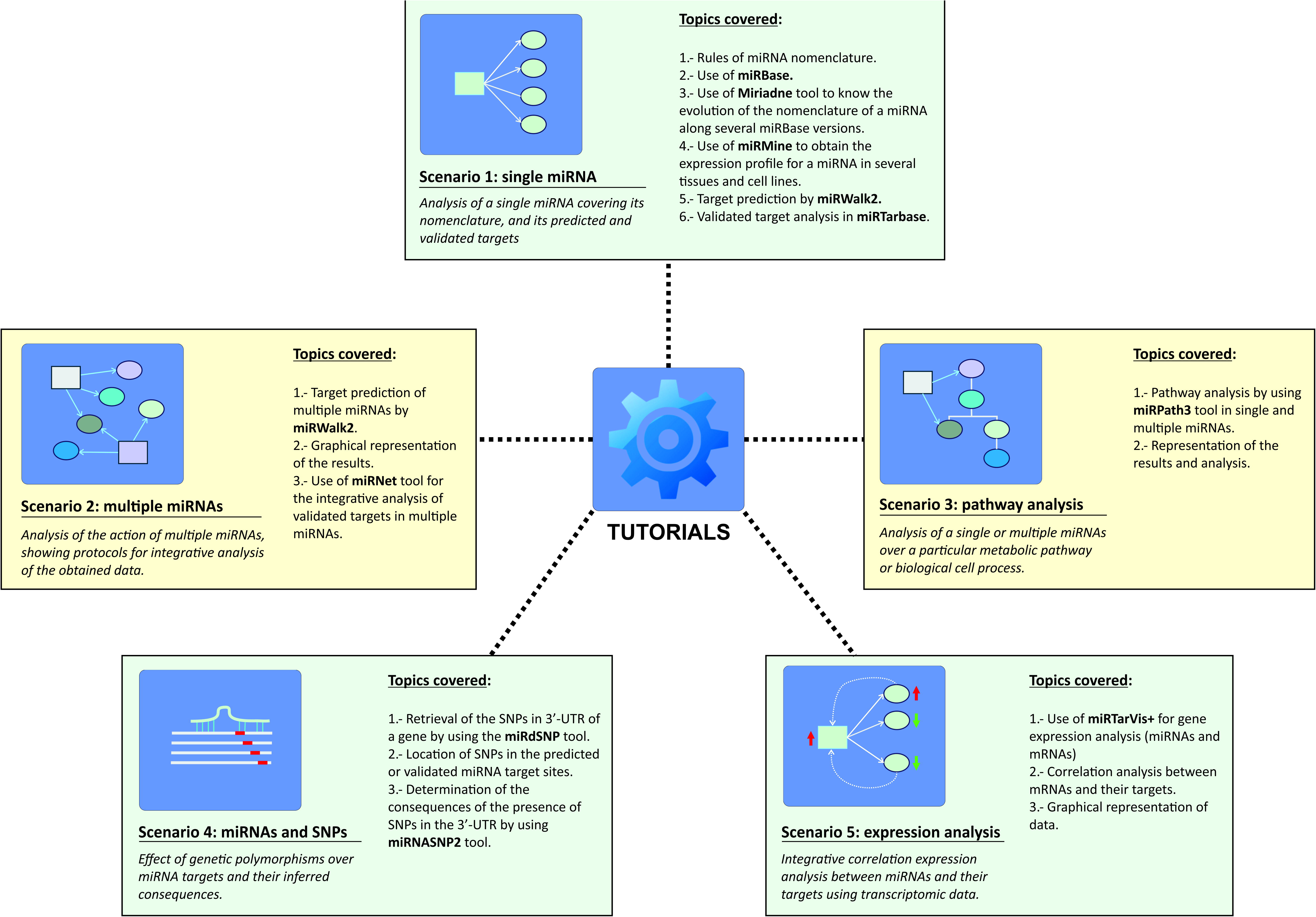
structure of the tutorials section of miRNAtools3 showing all the proposed working scenarios and the topics and computer applications covered in each of them.

### 4.1.- Scenario 1: single miRNA

The proposed scenario used a real example published by Asangani et al in 2008 [21] about an oncogenic miRNA, miR-21. The students are guided along the tutorial starting by simple steps such as retrieval of relevant data about the selected miRNA in miRBase [22, 23], and learn about proper miRNA nomenclature. The tutorial also refers about how to convert old-style miRNA nomenclature used in old releases of miRBase into the actual nomenclature which is use from miRBase v.18. In order to facilitate the nomenclature inter-conversion between miRBase versions, the tutorial explains how to utilize web-tools as Miriadne to follow the evolution of different nomenclatures for miR-21 along all miRBase versions [24]. This step is particularly relevant to conjugate the information about a particular miRNA that can be found in recent and old literature.

Following the tutorial the students will also learn how to retrieve expression data for the selected miRNA in different human cells and tissues by the use of miRmine, a recent database which compiles expression profile for miRNA in human tissues [15]. The final part of the tutorial comprises the retrieval of information about predicted and validated targets for the selected miRNA, hsa-miR-21-5p. For the prediction of miRNA targets the miRNAtools3 users are encouraged to use multiple predictors such as miRWalk2, since these applications are able to simultaneously interrogate several target prediction algorithms, displaying the results in a graphical manner and allowing the user to select those targets predicted by a higher number of algorithms [10]. For the retrieval of validated targets, the users are guided into the use of miRTarBase, a recently updated database of validated miRNA targets [9].

### 4.2.- Scenario 2: multiple miRNAs

For analyzing the simultaneous action of several miRNAs we proposed to study the targets of the miR-17-92a cluster, which is known to be overexpressed in some human tumors as lymphomas and involved in cell proliferation [25]. The main objective of this tutorial is to obtain information about the putative targets for the components of the miR-17-92a cluster, highlighting which of those targets could be more physiologically relevant. For this purpose we proposed to employ the strategy already described elsewhere [26], based on a circular layout graphical representation of the miRNA-target relationships that will allow to determine which mRNAs are simultaneously targeted by a higher number of miRNAs and which miRNAs are able to regulate more targets. We always stimulated the students to employ graphical representations for depicting the results obtained from the target prediction algorithms, since this kind of representations are easier to interpret and collect more information than the tabular formats. The tutorial explain how to construct this interaction networks by using a manual approach in the case of predicted targets or an automatic application for validated targets as MirNet [11].

### 4.3.- Scenario 3: pathway analysis

In our teaching experience, the students often get more stimulated when they can infer some physiological consequences of their data. This working scenario was designed with the objective of making some functional sense of the target prediction algorithms for a group of miRNAs. As in Scenario 2, we employed as a working hypothesis the human miR-17-92a cluster, but in this case the tutorial will guide the user in the prediction of the putative cellular pathways regulated by the six-membered miR-17-92a cluster. Among the existing web-tools for performing miRNA pathway analysis we selected a module of the DIANA-microT prediction suite, DIANA-miRPath v3.0, mainly because of its easy-to-use web interface and graphical output [16]. The tutorial guide the user to perform a pathway and GO-term enrichment analysis of the predicted and validated targets of the miR-17-92a cluster members by using miRPath3, and to produce graphical representations of the results. The tutorial explains also how to apply filters to the target information by relevance scores and the meaning of these scores in the context of a cellular pathway, guiding the users for a productive and meaningful interpretation of the obtained results.

### 4.4.- Scenario 4: single nucleotide polymorphisms and miRNA binding sites

This scenario is an interesting example of how the miRNA action can be considered within the field of pharmacogenomics. The proposed scenario is supported by a Genomic Wide Association Study (GWAS) published by Zuo and coworkers [27], which described some genetic variants in the gene encoding for the cannabinoid receptor (CNR1) that could be related with increased risk for cocaine dependency. In fact, one of these variantes named rs806368 is located in the 3’-UTR of CNR1 gene. The tutorial describes how to find information about the location of this SNP variant and the influence that this variant can eventually have in the putative binding of specific miRNAs. During the tutorial, the user is trained to use two databases: miRdSNP [28], a database of human disease-associated SNPs and miRNA targets and miRNASNP2 [29], a compilation of genotypic variants extracted from GWAS studies and related with miRNA binding sites. At the end of the tutorial the user should be able to analyze the effect of a particular genetic variant over the binding of a miRNA, by the calculation of the free energy of the miRNA-mRNA hybrid in the presence and in the absence of the genetic variant.

### 4.5.- Scenario 5: miRNA-mRNA expression correlation analysis

The biological action of a particular miRNA depends on the simultaneous existence of its cognate mRNA target(s) in the same cellular context. The characterization of the biological roles of miRNAs is often complex since cells are extremely redundant. Frequently, transcriptomic analysis is used as starting point to determine the relevance of action of miRNAs over the cell metabolism [30]. This tutorial describes how to use gene expression data to characterize the putative action of a group of miRNAs by using a correlation analysis. In a biological system an inverse correlation of expression levels between a miRNA and its cognate target is expected, however this effect could not be evident at the levels of mRNA transcript depending on the specific characteristics of the considered mRNA [31, 32]. In this tutorial we describe the use of a web-based tool named MirTarVis+ [14, 33], which is able to perform such kind of correlation analysis depicting the results in a graphical manner. The tutorial is based on real expression data collected by microarray analysis in tumor and normal cells. The corresponding data files are supplied to the user, and the specific format for input files in MirTarVis+ explained. The tutorial follows the logical flowchart of MirTarVis+, starting with data filtering and the inverse correlation analysis of miRNAs and their targets, and finishing with a graphical representation of the results that allow the user to determine which miRNAs and targets could be more relevant for this particular biological scenario.

## 5.- Conclusions and further developments

Advanced teaching initiatives for post-graduate students are essential for their integration in the scientific culture and methods, especially in those areas where there is rapid accumulation of scientific knowledge. We have designed a flexible web-tool specifically devoted for teaching about miRNA function by using only web-based applications that fulfilled an unmet need in this field of scientific knowledge. Our web-tool has been extensively improved and polished by its inclusion in tutorials given to students within workshops and courses dedicated to the study of miRNAs and other ncRNAs that have been taught in our institution and abroad.

Further developments of miRNAtools will be ensured by the intrinsic flexibility of the web design. The expansion of the number of proposed scenarios for teaching will take into account the valuable input received from the students and users of the webpage. Moreover, we are working in similar web-repositories for the study of lncRNAs and circRNAs that will be combined together with miRNAtools.

## ACKNOWLEDGEMENTS

The author wish to acknowledge the contribution of all the students involved in the training courses organized in our Institution and abroad, which have been essential to the improvement and development of miRNAtools3. We also acknowledge the suggestions coming from members of social networks that have been enthusiastically following every update of miRNAtools webpage. Finally, we are in debt with our colleagues from the UNESP, São Paulo, Brazil and specially to Prof. Ewa Stepien from the Jagiellonian University in Krakow, Poland, for their constructive discussions and improvements which have been implemented into every version of miRNAtools and for being hosts of our external editions of training courses about miRNAs and ncRNAs in cell biology and human disease.

